# Attenuated Dengue Virus PV001-DV Induces Oncolytic Cell Death and Potent Anti-Tumor Immunity

**DOI:** 10.1101/2022.07.05.498884

**Authors:** Josef W. Goldufsky, Preston Daniels, Michael D. Williams, Kajal Gupta, Bruce Lyday, Tony Chen, Geeta Singh, Andrew Zloza, Amanda L. Marzo

## Abstract

Viral therapies developed for cancer treatment have classically prioritized direct oncolytic effects over their immune activating properties. However, recent clinical insights have challenged this longstanding prioritization and have shifted the focus to more immune-based mechanisms. Through the potential utilization of novel, inherently immune-stimulating, oncotropic viruses there is a therapeutic opportunity to improve anti-tumor outcomes through virus-mediated immune activation. PV001-DV, is an attenuated strain of Dengue virus (DEN-1 #45AZ5) with a favorable clinical safety profile that also maintains the potent immune stimulatory properties known of Dengue virus. In this study, we examined the anti-tumor effects of PV001-DV as a potential novel cancer immunotherapy. In vitro assays demonstrated that PV001-DV possesses the ability to directly kill human melanoma cells lines as well as patient melanoma tissue ex vivo. Importantly, further in vitro work demonstrated that, when patient peripheral blood mononuclear cells (PBMCs) were exposed to PV001-DV, a substantial induction in production of apoptotic factors and immunostimulatory cytokines was detected. When tumor cells were cultured with the resulting soluble mediators from these PBMCs, rapid cell death of melanoma and breast cancer cell lines was observed. The direct tumor-killing and immune-mediated tumor cytotoxicity facilitated by PV001-DV contributes support of its upcoming clinical evaluation in patients with advanced melanoma who have failed prior therapy (NCT03989895).

## Introduction

The success of immunotherapy research and translation over the last 10 years has improved the quality-of-life and survival of patients with cancer. However, further improvements must be made to increase the proportion of patients that achieve long-term outcomes. Such improvement may be possible via the utilization of viruses that demonstrate inherent capabilities in facilitating anti-tumor responses. The class of oncolytic viruses has best demonstrated this clinical benefit, most notably with the approval of TVEC for the treatment of metastatic melanoma. [1p] Oncolytic viruses inherently possess the ability to infect and lyse tumor cells, which can occur in an immunogenic manner, thus releasing tumor antigens to be scavenged by antigen-presenting dendritic cells (DCs) and generating tumor-specific T cell responses. Furthermore, many oncolytic viruses express immuno-stimulatory transgenes to reshape the tumor microenvironment to increase this therapeutic effect. [2p, 3p] However, few viruses with inherent potent immune activating properties have been investigated as viral-based therapies for cancer as prevailing sentiment has been that anti-viral immunity may be a detriment to the oncolytic effect. [4p, 60p] Recent clinical observations, however, have noted improved outcomes in patients exhibiting anti-viral immune responses. [5p, 6p, 7p, 8p] The potential of anti-viral immune responses to facilitate improved anti-tumor responses creates an opportunity to develop a new class of virotherapies with more robust immune-activating properties and differentiated anti-tumor mechanisms.

Many viruses have an increased capacity to infect and kill tumor cells compared with normal, non-transformed cells [9p]. This phenomenon, in part, has been attributed to several factors, including increases in growth factor receptors on tumor cells that co-serve as viral entry receptors, as well as the propensity of tumor cells to disable their interferon (IFN) gene signaling pathways as an immune evasion mechanism. Without an intact IFN network, viral replication can preferentially proceed at a rapid rate within tumors, often leading to tumor cell death by a cytopathic effect [10p]. While many oncolytic viruses are large DNA viruses that have been modified with transgenes to make them selective for tumor cells and have increased immune stimulation properties, unmodified RNA viruses such as the Coxsackie virus, Newcastle virus, and Reovirus have also been evaluated as oncolytic therapies. [11p, 12p] The capacity of wild-type RNA viruses to achieve high viremia and tumor infection rates, well suits them as effective immune activators triggering more acute immune responses that facilitate more potent cell killing potential.

Dengue virus (DV) innately possess favorable properties as a novel virotherapy in the age of immunotherapy. Expression of the heparan sulfate proteoglycan (HSPG) receptor and Dendritic Cell-Specific Intercellular adhesion molecule-3-Grabbing Non-integrin (DC-SIGN) on many tumor subtypes allows effective binding of and cell entry into tumor cells [13p]. In prior pre-clinical investigations, dengue virus has demonstrated the capacity to kill neuroblastoma, hepatic, and leukemia tumor cells via super-oxides, arachidonic acid, and endoplasmic stress. [14p, 15p, 16p] In addition to this observation, upon human infection with dengue virus, pro-inflammatory and Th_1_ cytokines are produced, which leads to vigorous activation of both the innate and adaptive immune systems. [17p] DV infection of monocytes, DCs and other PBMCs causes up-regulation of genes associated with both innate and adaptive responses. [18p, 19p, 20p] These gene products are secreted as chemokines and cytokines, including those capable of causing cell death via apoptosis factors binding to specific receptors on the cell membrane. [21p, 22p] This commonly observed clinical immune response with Dengue virus possesses many characteristic overlaps with the desired immune response in effective anti-tumor responses, giving strong rationale for investigation of this virus as a cancer viro-immunotherapy.

For this study we used PV001-DV, a live-attenuated strain of Dengue Virus (DEN-1 #45AZ5). PV001-DV has completed investigation in a phase 1 challenge study (NCT02372175) and resulted in uncomplicated dengue illness in healthy volunteers [23p]. The results of this phase 1 study demonstrated effective viral replication and immune activation while maintaining an appropriate safety profile for clinical development as a novel immunotherapy in oncology. In this study, we examined the novel anti-tumor effects of PV001-DV as well as the ability of this virus to facilitate a potent anti-tumor immune response.

## Methods

### Virus

Dengue Virus #45AZ5 (PV001-DV) was obtained by Primevax Immuno-Oncology, Inc., from the Walter Reed Army Institute of Research (WRAIR) in Silver Spring, MD., under license with the US Army Medical Materials Development Activity, Ft. Detrick, MD. PV001-DV was shipped as a lyophilized powder and was reconstituted with RPMI 1640 media to 650,000 pfu/1000 μL.

### Tumor Cells

The human melanoma cell lines A2058, SK-MEL-2, SK-MEL-5, and SK-MEL-28 as well as human breast cancer cell line MDA-MB-231 were obtained from the American Type Culture Collection (ATCC). Cells were thawed, centrifuged, and expanded in recommended culture media, as directed by the manufacturer, and then, cryopreserved for later expansion and further use.

### Cytopathic Effect in vitro

Frozen cell lines were thawed and cultured overnight in recommended complete media as per ATCC *Handling Information*. At 70-80% confluence, cells were removed from flasks with Trypsin-EDTA chelating agent, counted, and plated in 24-well tissue culture treated plates in triplicate at 30,000 cells per well. Cells were incubated at 37°C and 5% CO_2_ for up to 6 days, as described in the Results section, and imaged or utilized for flow cytometry studies. During incubation, cells were treated with PV001-DV at the pfu (e.g., 0, 1,000, 5,000 and 25,000 pfu), or as indicated for each experiment.

### Imaging

Cells in 96-well plates were imaged using a Keyence BZ-X800 microscope. Images are shown at the magnification indicated for each respective study within the Results section.

### Flow Cytometry

Cells were stained with propidium iodide (PI, Millipore Sigma, P4864-10ML), ApoTracker (BioLegend, 427403), Live/Dead Fixable Aqua (ThermoFisher, 501121526) or antibodies to DC-SIGN (BD Biosciences, 564127), Syndecan-2 (ThermoFisher, 25-1389-42) and PD-L1 (BD Biosciences, 740426), as described for each study in the Results section. Flow cytometry was conducted using a BD LSR2 cytometer. Analysis of flow cytometric data was performed using FlowJo v.10. Gating strategies are listed for each respective study within the Results section.

### Patient Tumor and PBMCs

A patient with cutaneous melanoma and a patient with breast cancer signed research informed consents. Tumor tissue and/or peripheral blood were obtained under IRB approval by the Rush University Biorepository Core (RUBC), which provides researchers access to clinical de-identified biospecimens.

### Tumor Tissue ex vivo

The melanoma tissue was bisected and each ∼2 mm x 2 mm piece was placed in a 96-well tissue culture treated, flat-bottom plate with complete RPMI 1640 media and incubated at 37°C and 5% CO_2_. During incubation the tumor tissue was treated with PV001-DV at 65,000 pfu or media control and then imaged at 24-hour intervals over 72 hours. Further, at 72 hours, the tissue was mechanically dissociated using a gentleMACS Octo Dissociator (Miltenyi Biotec), enzymatically dissociated in HBSS containing 1 mg/mL type IV collagenase from Clostridium histolyticum (Sigma-Aldrich) and 40 μg/mL DNAse I from bovine pancreas (Sigma-Aldrich) at 37°C for 30 minutes with constant rocking, and then mechanically dissociated again, as previously described [64p]. A 70-μm filter was utilized subsequently to yield a single-cell suspension. Cells were then analyzed by for flow cytometry, as indicated within the Results section.

### PBMCs ex vivo

PBMCs were isolated from melanoma and breast cancer patient peripheral blood (∼10 ml) by Ficoll gradient (Lymphoprep) centrifugation, washed in PBS, and plated at 100,000 cells/well in triplicate wells of a 96-well plate in complete RPMI 1640 media. PBMCs were incubated for 72 hours with PV001-DV added to the wells at a ratio of 2 pfu of virus per PBMC (i.e., MOI = 2) or with RPMI 1640 vehicle control. Well contents were centrifuged at 1400 rpms for 10 minutes and supernatants were obtained. Cell-free supernatants were then filtered using Whatman 20 nM Anotop sterile syringe filters to remove virus particles. Cell-free, virus-free supernatants were then cultured with A2058, SK-MEL-5, and SK-MEL-28 cells for 24 hours. Tumor cell lines were then imaged and used in flow cytometric analysis.

### Soluble mediator identification

PBMC-derived soluble mediators were identified within the cell-free, virus-free supernatants (described in the *PBMCs ex vivo* methods; 50 μl each) using the Immune Monitoring 65-Plex Human ProcartaPlex Panel (ThermoFisher, Catalog number: EPX650-16500-901), as per manufacturer instructions. The mediators (biomarkers) concentrations were calculated using a 5-parametric curve fit using xPONENT, version 4.03 (Luminex) in a blinded fashion with measurement performed with the FlexMAP 3D system (Luminex).

## Results

### PV001-DV induces apoptotic cell death of human melanoma cell lines

Dengue virus has been demonstrated to have tumor killing ability in previous pre-clinical investigations. [14p, 15p, 16p] To confirm these observations using PV001-DV, we examined the direct tumor killing efficiency of this strain in commonly utilized human melanoma cell lines. Initially, A2058 melanoma cells were co-cultured with PV001-DV **(Figure 1A)**. Preliminary visual observations at 3 days indicated increased numbers of dead cells and reduced confluency in the cells cultured with 25,000 pfu of PV001-DV compared to the 0 pfu control **(Figure 1B)**. Final analysis of cultured cells at 6 days by propidium iodide and ApoTracker confirmed that PV001-DV kills A2058 melanoma cells and reduces their viability through facilitating tumor cell apoptosis and necrosis **(Figure 1C, 1D)**. To ensure that this direct anti-tumor effect was not limited to a single cell line, additional melanoma cell lines (SK-MEL-2 and SK-MEL-28) were cultured with PV001-DV under similar conditions **(Figure 1E)**. These cells also underwent cell death within 2 days as detected by Aqua Live/Dead viability staining **(Figure 1F,G)**. Ultimately, these results confirm that PV001-DV possesses direct anti-tumor activity against human melanoma cells.

**Figure 1.**
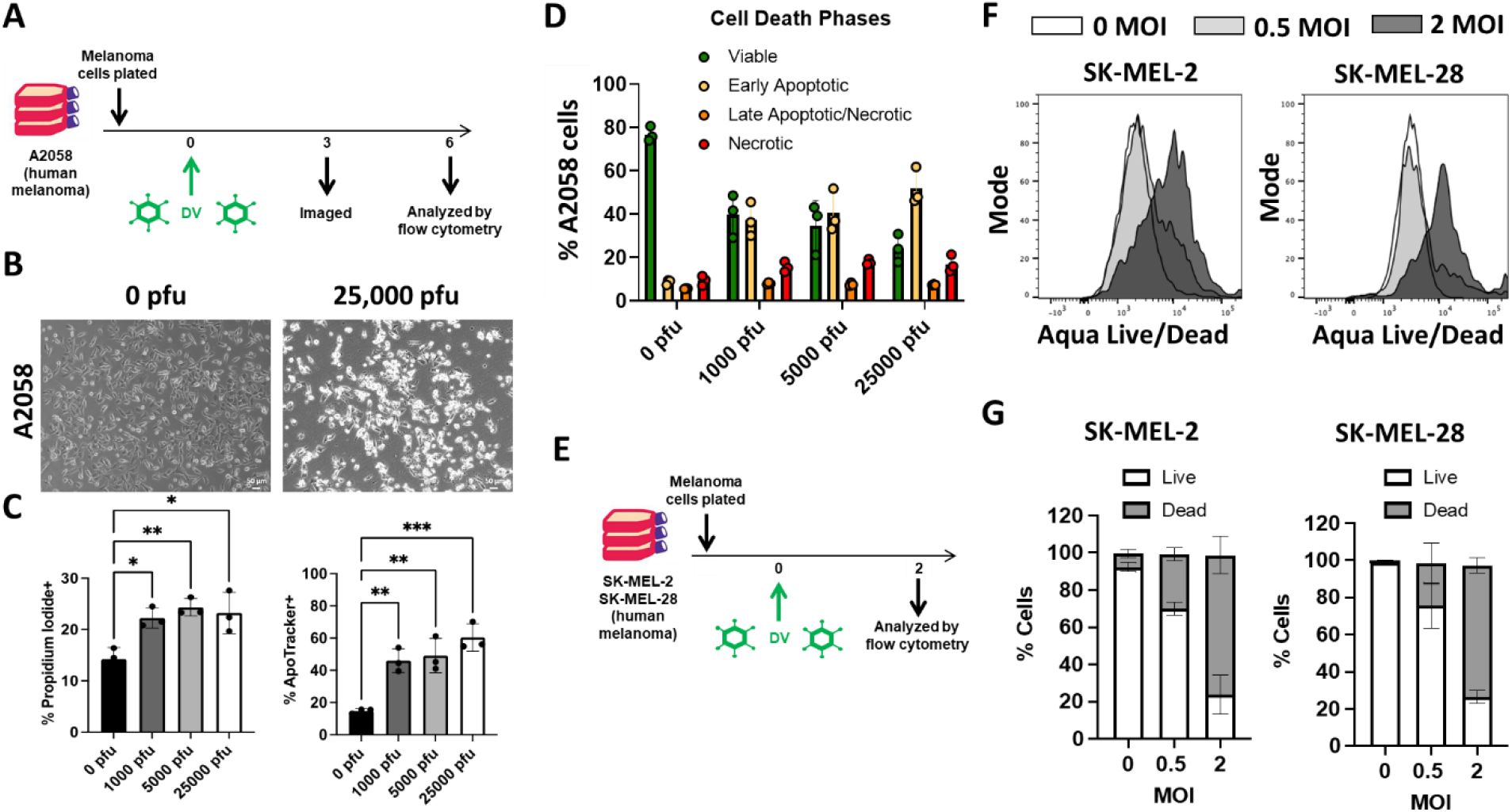
PV001-DV exhibits direct oncolytic effect on human melanoma cell lines. **(A)** Schematic of A2058 human melanoma cells incubated with PV001-DV (DV) and measured for viability. **(B)** Photomicrograph of cells at 10X magnification from (A) at 3 days of incubation. **(C)** Percent Propidium Iodide (PI) and percent ApoTracker staining of cells from (A) at 6 days. *P <0.05, **P <0.01, ***P <0.001, One-way ANOVA with Tukey correction. **(D)** Division of cells from (A) into viable, early apoptotic, late apoptotic/necrotic, and necrotic based on combination PI and ApoTracker staining. **(E)** Schematic of SK-MEL-2 and SK-MEL-28 melanoma cells incubated with PV001-DV (DV) and measured for viability. **(F)** Flow cytometry histograms of viability via Aqua Live/Dead staining from experiment in (E). **(G)** Bar graphs of viability from experiment in (E).

### PV001-DV direct cell killing of melanoma patient tumor

Due to differences in morphology and the impact of extend passages, cell lines can exhibit different sensitivities than primary tumors. This is particularly due to the three-dimensional nature of tumors versus in vitro cell lines. [24p] To further substantiate the clinical relevance of the direct tumor cell-killing potential of PV001-DV observed in Figure 1, a primary resection sample of advanced cutaneous melanoma was obtained and cultured with PV001-DV. When the sample was examined visually and by flow cytometry after 3 days, substantial tumor cell killing was observed **(Figure 2A-C)**. Specifically, cells decellularizing from the tissue demonstrated expected melanin (dark/black) pigment staining in the control, while those cells decellularizing in the PV001-DV-treated sample were devoid of pigment, thus demonstrating compromised membranes indicative of cell death progression **(Figure 2B)**. Further, after mechanical and enzymatic tumor tissue dissociation, the resultant cells demonstrated substantial tumor killing in the PV001-DV-treated group after staining with Aqua Live/Dead **(Figure 2B)**. These results further confirm the direct anti-tumor potential of PV001-DV and its ability to infect and kill tumor cells.

**Figure 2.**
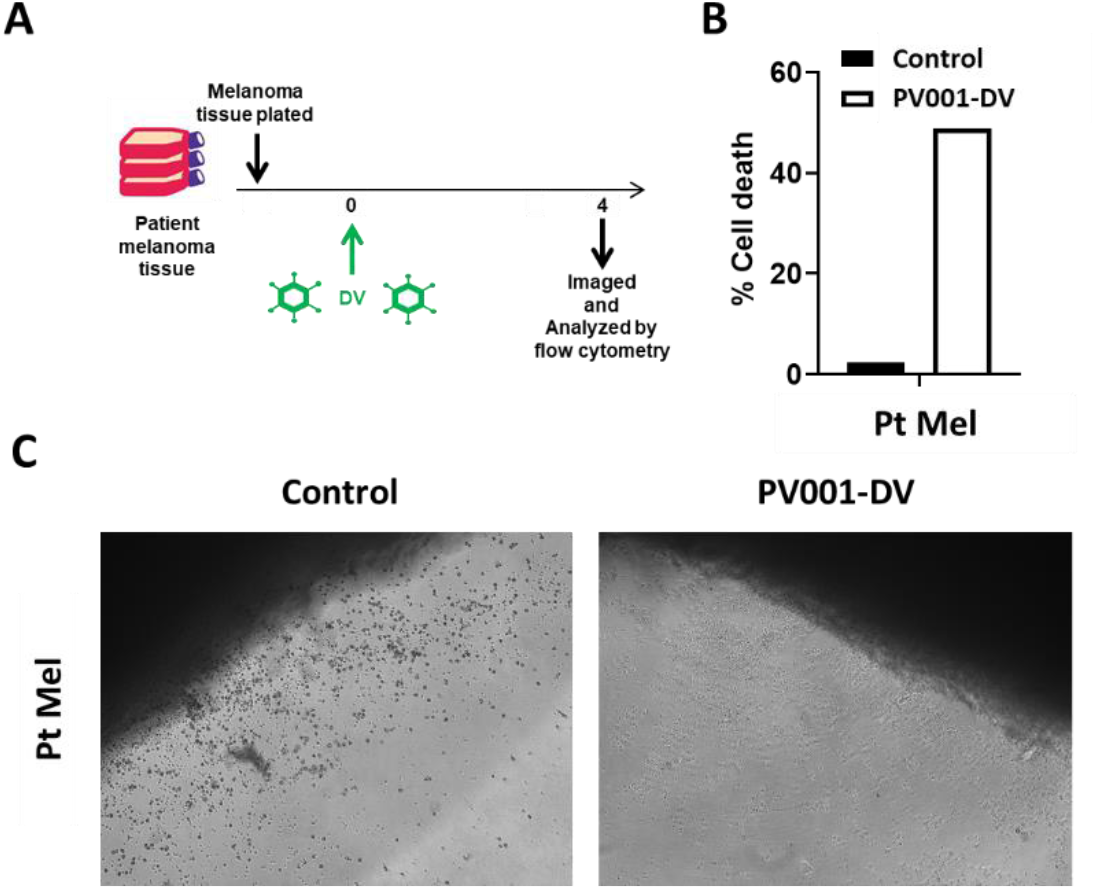
PV001-DV exhibits direct oncolytic effect on patient-derived melanoma tumor. **(A)** Schematic of patient-derived melanoma sample incubated with PV001-DV (DV) and measured for viability. **(B)** Percent tumor cell death as measured by Aqua Live/Dead staining via flow cytometry from experiment in (A). **(C)** Photomicrograph at 10X magnification at 4 days of incubation from experiment in (A).

### Cancer patient-derived PBMCs exposed to PV001-DV secrete soluble factors that exert potent anti-tumor effects

Typical Dengue virus infection, though often mild, results in an inflammatory febrile disease characterized by a predictable course of acute immune activation. This immune response is often robust and systemically engages a variety of important cell types. [17p, 18p, 19p, 20p] Due to the mediators of acute anti-viral responses having many overlaps with the mediators of an anti-tumor response, it is believed that this response can be co-opted for anti-tumor activity. [7p, 8p] To examine this phenomenon, we exposed melanoma patient-derived PBMCs to PV001-DV for 3 days at which point cell-free, virus-free, supernatant was isolated and co-cultured with human melanoma cell lines **(Figure 3A)**. At 1 day the A2058 and SK-MEL-5 cells with uninfected PBMC supernatants showed characteristic elongated morphology of proliferating tumor cells, while the wells with DV-infected PBMC supernatants showed markedly fewer cells, with large amounts of the visual field devoid of cells, or with many rounded cells showing evidence of apoptosis **(Figure 3B)**. Similarly, by Aqua Dead/Live staining, A2058 and SK-MEL-5 cells exhibited greater killing of tumor cells exposed to PV001-DV PBMC supernatant compared to controls **(Figure 3C)**. To determine whether this supernatant effect could be reproduced by utilization of PBMCs from a patient with a different cancer type, we conducted a study similar as for the melanoma patient-derived PBMCs, however, this time using breast cancer patient-derived PBMCs. For MDA-MB-231 breast cancer cells, the killing of tumor cells exposed to PV001-DV PBMC supernatant was substantially increased compared to controls **(Figure 3D)**. Interestingly, breast cancer patient-derived PV001-DV PBMC supernatant also increased killing of melanoma cell lines, A2058 and SK-MEL-5, exhibiting a cross-tumor effect of the supernatant **(Figure 3D)**. Overall, these results demonstrate the ability of the anti-viral immune response to PV001-DV to generate extracellular mediators (cell-free and virus-free) from cancer patient-derived PBMCs that result in human cancer cell death within a short period of time. Importantly, these effects were observed across different cell lines and tumor types, showing robust and broad anti-tumor immune activation.

**Figure 3.**
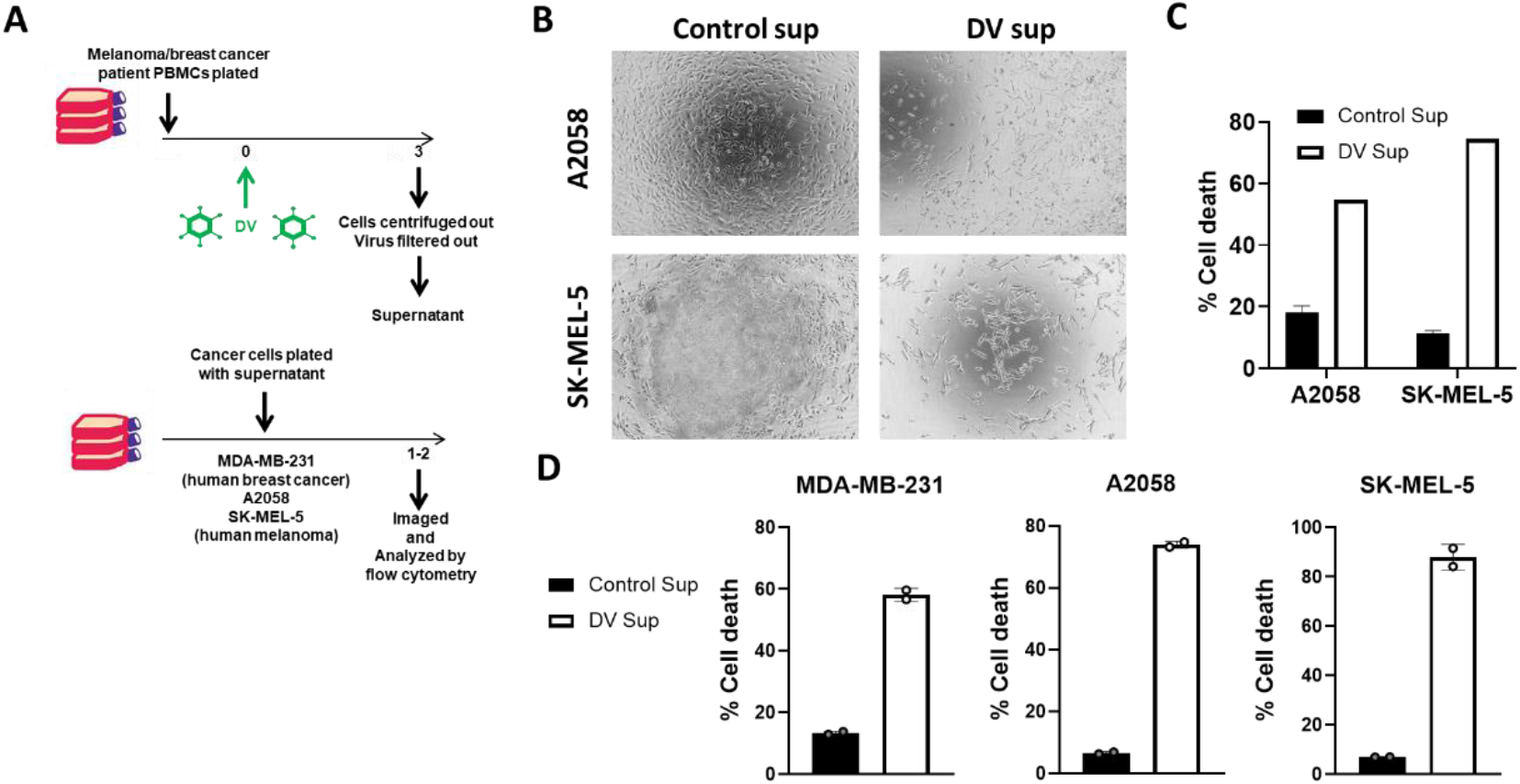
PBMCs treated with PV001-DV produce cytotoxic soluble mediators. **(A, top)** Schematic of melanoma and breast cancer patient PBMCs incubated with PV001-DV (DV) for 3 days. Cells well removed by centrifugation and virus was removed by filtration. **(A, bottom)** Schematic of resultant supernatant incubated with A2058 and SK-MEL5 melanoma or MDA-MB-231 breast cancer cells. **(B)** Photomicrograph of cells at 10X magnification from (A) at 1 day of incubation. **(C)** Viability of melanoma cells measured with Aqua Live/Dead from experiment in (A) after incubation with melanoma patient PBMC-derived supernatant. **(D)** Viability of breast cancer and melanoma cells measured with Aqua Live/Dead from experiment in (A) after incubation with breast cancer patient PBMC-derived supernatant.

### PV001-DV PBMC supernatant increases known dengue virus binding ligands and immune checkpoint receptor, PD-L1

Epigenetic and phenotypic changes can occur in cells when exposed to inflammatory conditions. This is true of cancer cells as well. [27p, 28p, 29p] We set out to examine the expression changes that may facilitate variable sensitivity to continued propagation of PV001-DV during an active immune response. To evaluate this, viable cancer cells remaining after exposure to PV001-DV PBMC supernatant were examined for changes in expression of known ligands for Dengue virus. Dengue virus utilizes DC-SIGN and heparan sulfate to enter host cells. [30p, 31p, 32p, 33p, 34p] Syndecan-1, is a modified heparan sulfate proteoglycan (HSPG) that DV uses to infect cells that is upregulated on tumor cells and increases the migratory potential of melanoma cells. [31p, 35p] Soluble mediators secreted from PBMCs exposed to PV001-DV increased the levels of both primary ligands for Dengue virus, DC-SIGN and Syndecan-1 **(Figure 4A,B)**. Importantly, PD-L1 expression is well-characterized to be elevated in some inflammatory microenvironments. [36p, 37p] While typically a poor prognostic factor, PD-L1 upregulation in response to an inflammatory microenvironments is a relatively favorable biomarker for response to anti-PD-1 therapy [38p]. Therefore, we examined its expression and found it to likewise be increased following co-culture with PV001-DV PBMC supernatant **(Figure 4A,B)**. These results suggest that enhanced PV001-DV tumor binding and infection rates could occur during the immune response further accelerating direct cytotoxic effects of PV001-DV. This combination of these mechanisms could lead to synergistic increases in tumor cell killing in vivo. Furthermore, this response could prime the immune response to facilitate improved outcomes with follow-on immune checkpoint inhibitor therapy.

**Figure 4.**
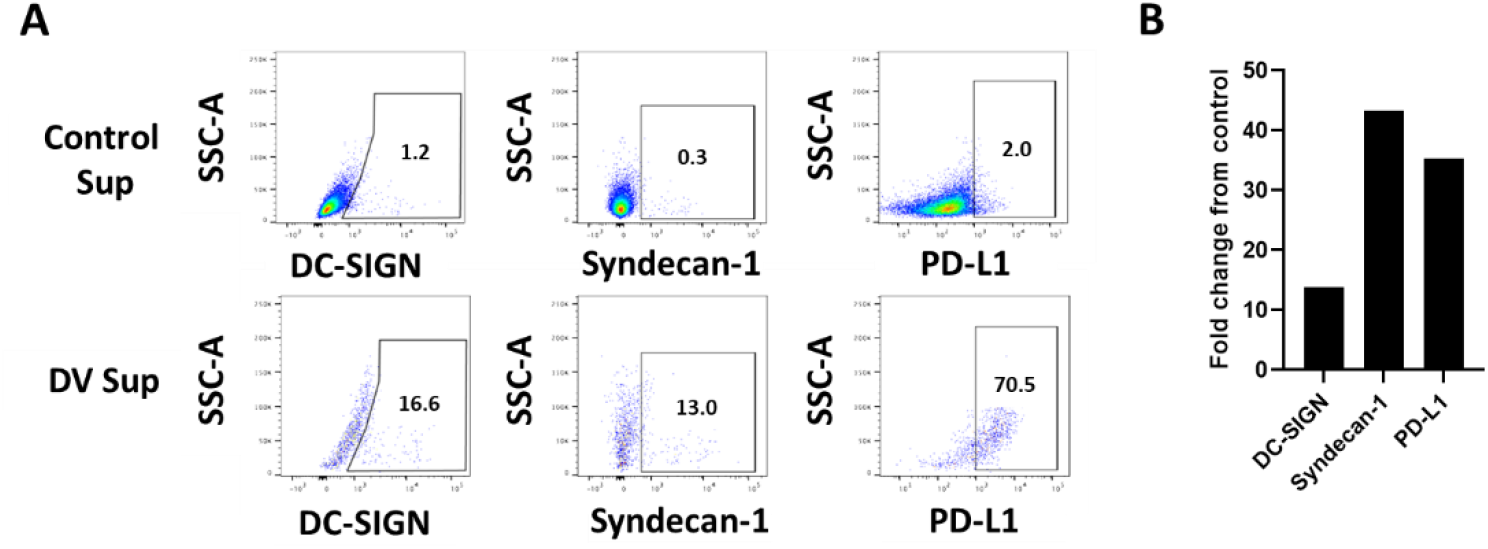
Supernatant from PBMCs cultured with Dengue virus increases surface receptors for Dengue virus cell binding and entry as well as immune checkpoint inhibition on tumor cells. **(A)** Flow plot analysis of MDA-MB-231 tumor cells by flow cytometry for expression of Dengue virus receptor ligands, DC-SIGN and Syndecan-1, as well as immune checkpoint receptor, PD-L1. **(B)** Fold change of receptors comparing levels from the PV001-DV supernatant treatment (DV sup) over media control (Control sup).

### PBMCs exposed to PV001-DV increase secretion of known apoptotic factors

Upon exposure to viruses, immune cells produce and secrete a variety of soluble factors that mediate the corresponding immune response and help facilitate the suppression of the virus, including inducing cell death in virally infected cells. [25p, 26p] To further examine the anti-tumor cytotoxic response observed, we ran a Luminex analysis to characterize the changes in the secretion of 65 immune-related soluble factors (cytokines, chemokines, and growth factors) by PBMCs following exposure to PV001-DV. Supernatants of respective cancer patient PBMCs not exposed to PV001-DV were used as a control. Relative values observed in secretion were plotted as a heat map (**Figure 5A-C**). Overall, a wide breadth of immune-related soluble factors were found increased in the supernatants from melanoma (55/65) and breast cancer (60/65) PBMCs exposed to PV001-DV over respective control supernatants (**Figure 5A**). Prior to conducting the assay, we identified a subset panel of factors known to mediate the induction of apoptosis in tumor cells and their additional corresponding immune activity **(Table 1)**. In the context of supernatants from melanoma patient PBMCs exposed to PV001-DV, an increase in 14/16 of the apoptotic factors secreted was observed over control supernatants **(Figure 5B)**. In supernatants from breast cancer patient PBMCs exposed to PV001-DV, increases in 15/16 of the apoptotic factors secreted were observed over control **(Figure 5B)**. These data demonstrate the potential of PV001-DV to activate a robust and broad immune response that can facilitate apoptotic activity in tumors as well as further downstream immune activation and response.

**Figure 5.**
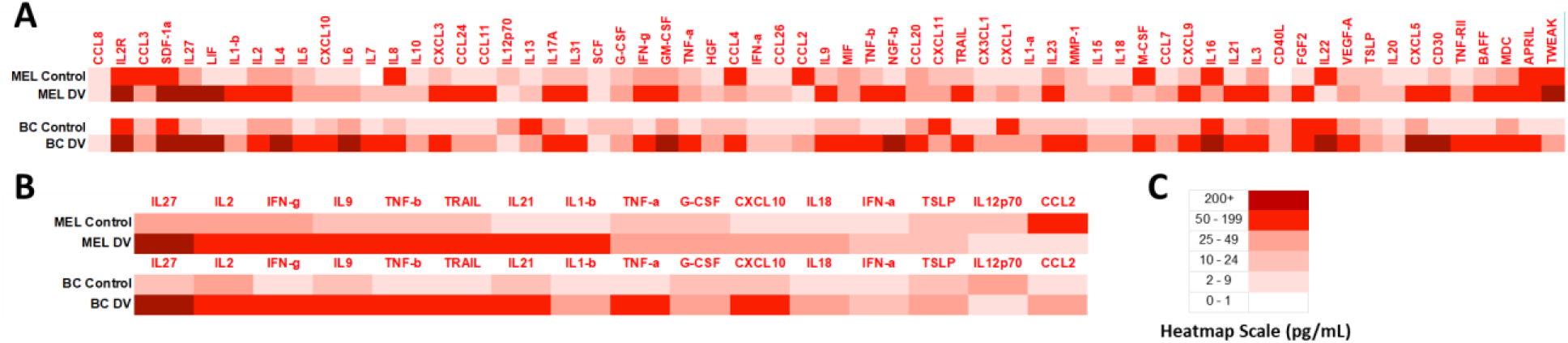
Increases in pro-apoptotic proteins secreted by PBMCs treated with PV001-DV. **(A)** Heatmap of soluble mediators measured from melanoma and breast cancer patient PBMC-derived supernatants via the Invitrogen Immune Monitoring 65-Plex Human ProcartaPlex Panel. Heatmap of an a priori-selected pro-apoptotic panel. **(C)** Legend of heat map ranges.

**Table 1.**

| Analyte | Immune Properties | Apoptotic Activity |
| --- | --- | --- |
| **Interleukin-27 (IL-27)** | IL-12 family cytokine, immune activation | Induces tumor cell apoptosis [39p] |
| **Interleukin-1 beta (IL-1β)** | Inflammatory cytokine, promotes fever | Induces tumor cell apoptosis [40p] |
| **Interleukin-2 (IL-2)** | Promotes T cell proliferation and effector function | Can induce apoptosis in tumor cells [41p] |
| **Chemokine ligand -10 (CXCL10)** | Recruitment of lymphocytes including T cells | Induces tumor cell apoptosis [42p] |
| **Interleukin-12 p70 (IL-12p70)** | Key cytokine for anti-tumor Th<sub>1</sub> responses | Induces tumor cell apoptosis [43p] |
| **Granulocyte-Colony Stimulation Factor (G-CSF)** | Promotes neutrophil development | Can induce apoptosis in tumor cells [44p] |
| **Interferon-Gamma (INF-γ)** | Critical cytokine for anti-tumor Th<sub>1</sub> responses | Key apoptosis factor for tumor cells [45p] |
| **Tumor-Necrosis Factor Alpha (TNF-α)** | Inflammatory cytokine, fever inducer, shock in high amounts | Key apoptosis factor for tumor cells [46p] |
| **Interferon alpha (INF-α)** | Key cytokine for antiviral and antitumor responses | Key apoptosis factor for tumor cells [47p] |
| **Chemokine ligand -2-MCP-1 (CCL2)** | Innate cell trafficking | Can induce apoptosis in tumor cells [48p] |
| **Interleukin-9 (IL-9)** | Activates T cells | Can induce apoptosis in tumor cells [49p] |
| **Tumor necrosis factor Beta (TNF-β)** | Key innate signaling cytokine | Can induce apoptosis in tumor cells [50p] |
| **Tumor Necrosis Factor Apoptosis Inducing Ligand (TRAIL)** | Attaches to Death Receptors DR3/4 on tumor cell surface | Key apoptosis factor for tumor cells [51p] |
| **Interleukin-18 (IL-18)** | IL-1 family proinflammatory cytokine | Can induce apoptosis in tumor cells [52p] |
| **Interleukin-21 (IL-21)** | Key T cell cytokine of IL-2 family, promotes T cell effector and memory functions, NK activity | Key apoptosis factor for tumor cells [53p] |
| **Thymic Stromal Lymphoprotein (TSLP)** | Activates dendritic cells | Can induce apoptosis in tumor cells [54p] |

## Discussion

PV001-DV is an attenuated strain of Dengue virus that is being developed for the treatment of solid tumors. In a phase 1 clinical study in healthy volunteers this strain was well-tolerated and manifested as an uncomplicated dengue illness. [23p] Dengue virus has inherent tropism for tumors with elevated expression of HSPG, DC-SIGN, and its other natural receptors. This allows for Dengue virus and PV001-DV to replicate within tumor cells, thus compromising their viability and leading to virus-induced cell death. [30p, 31p, 32p, 33p, 34p, 35p] Furthermore, dengue preferentially replicates within the tumor given the often-compromised and reduced host cell immune activity allowing for enhanced direct tumor cell killing and viral propagation to occur. [10p] Viral-induced cell death increases the danger-associated molecular pattern (DAMP) response as well as the release of neo-epitopes. This stimulates anti-tumor immune activity and allows for the generation of tumor-antigen specific response to the newly presented tumor epitopes [2p, 55p]. These are hallmarks of the approach of many oncolytic viruses used in clinical development and Dengue virus has preliminarily demonstrated comparable ability in facilitating these direct anti-tumor effects.

The original and longstanding oncolytic paradigm is to primarily exert anti-tumor effects through viral propagation and oncolysis. Immune effects have been a secondary mechanism with reliance on inherent immune activation through DAMP production during cell lysis. More recently, transgenes have been added to oncolytic viruses to amplify specific immune signals to facilitate defined immune mechanisms. The broader anti-viral immune response, however, has largely been considered a detriment to response due to limiting viral propagation. [2p, 3p, 4p, 56p, 60p] To this effect, investigation of oncolytic viruses has focused on viruses with the ability to infect and kill tumor cells by oncolysis, but only weakly stimulate effector T and NK lymphocytes. However, recent clinical studies have reported enhanced responses in patients who have demonstrated anti-viral immune responses challenging the long-standing paradigm of the preference of oncolytic viruses to avoid an anti-viral immune response. [5p, 6p, 7p, 8p]

New hypotheses have emerged around the benefit of an anti-viral response in facilitating an anti-tumor immune response attributing to the many overlaps in the responding immune subtypes. The ability of a virus to not only exert direct anti-tumor effects through oncolysis, but also to serve, as an anti-tumor immune adjuvant allows for the enhanced priming of an anti-tumor immune response [57p]. Furthermore, the generation of a primary acute immune response by a virus facilitates the complete immune response cycle. This robust activation accounts for stimulation of multiple immune subtypes and programs that are required for the anti-tumor immune response but may go unaddressed by the expression of single cytokine transgenes delivered by classic modified oncolytic viruses. [18p, 19p, 20p, 58p]

This robust immune activation by an immunostimulatory virus can be co-opted for priming local and systemic anti-tumor immune responses. A major hurdle in cancer therapy is the influence of an immunosuppressive tumor microenvironment on reducing the functional capacity of an anti-tumor immune response. This immunosuppression can have profound effects on even the most potent immunotherapies in our arsenal. A prognostic factor for improved outcomes following CAR-T cell or anti-PD-1 therapy is the amount of immune suppression in the tumor microenvironment [59p, 61p]. Addressing this challenge by “heating up” the tumor microenvironment with a virus that retains oncolytic potential but also induces potent inflammatory immune activity would be a welcome addition to the clinical arsenal of immunotherapies for cancer and provides rationale for potential combination therapies.

Facilitating the stimulation of necessary immune subtypes is one potential benefit of using more inherently immunostimulatory viruses, but the downstream effects of this can have further anti-tumor impact. When immune cells are stimulated with a potent virus like Dengue, they secrete immunomodulatory cytokines that amplifies the immune response by additional immune cell subtypes. Through this cascade, we have demonstrated that immune cell subsets begin the secretion of apoptotic factors that kill cells with compromised integrity. This response is intended to kill virally infected cells to quell viral propagation, but also has the unique ability to kill tumor cells as well. [25p, 26p] Dengue virus appears to be potentially potent in this function and have unique capabilities in stimulating this response. [62p] The ability of an oncolytic virus to use apoptosis factors to kill tumors in a manner independent, yet potentially synergistic with oncolysis and enhanced effector cell function, provides a novel approach to improve clinical responses.

PV001-DV is differentiated from other oncolytic viruses in that it processes inherent potent immune activation that enhances cytotoxic immune activation. However, it is also differentiated from other immunostimulatory viruses given its ability to direct tumor cell killing. Furthermore, viruses with immune-activating properties also tend to be more virulent, limiting their potential development as therapeutic candidates. Dengue virus has a favorable safety profile compared to other potent viruses with a death rate of 0.025%, with most of these deaths occurring as amplified heterologous secondary infections in areas without modern supportive medical care [63p]. By use of the live but attenuated strain, PV001-DV, in areas with low risk of secondary heterologous infection, safety will be dramatically improved from this already favorable figure. This is further supported by prior clinical characterization of this well-tolerated strain and backs the further clinical development of this strain as a novel cancer immunotherapy. [23p]

In conclusion, PV001-DV specifically has the capacity to directly infect and kill tumor cells. Furthermore, PV001-DV has immune potentiating properties that facilitates increases in the production of cytokines that enhance effector cell function and indirect cytotoxic factor production. These observations provide a strong basis for support of the potential as PV001-DV as a cancer immunotherapy and further support forthcoming clinical investigation in advanced melanoma (NCT03989895).

